# Video-rate Mid-infrared Photothermal Imaging by Single Pulse Photothermal Detection per Pixel

**DOI:** 10.1101/2023.02.27.530116

**Authors:** Jiaze Yin, Meng Zhang, Yuying Tan, Zhongyue Guo, Hongjian He, Lu Lan, Ji-Xin Cheng

## Abstract

By optically sensing the mid-infrared absorption induced photothermal effect, midinfrared photothermal (MIP) microscope enables super-resolution IR imaging and scrutinizing of biological systems in an aqueous environment. However, the speed of current lock-in based sample-scanning MIP system is limited to 1.0 millisecond or longer per pixel, which is insufficient for capturing dynamics inside living systems. Here, we report a single pulse laserscanning MIP microscope that dramatically increases the imaging speed by three orders of magnitude. We harness a lock-in free demodulation scheme which uses high-speed digitization to resolve single IR pulse induced contrast at nanosecond time scale. To realize single pulse photothermal detection at each pixel, we employ two sets of galvo mirrors for synchronized scanning of mid-infrared and probe beams to achieve an imaging line rate over 2 kHz. With video-rate imaging capability, we observed two types of distinct dynamics of lipids in living cells. Furthermore, by hyperspectral imaging, we chemically dissected a single cell wall at nanometer scale. Finally, with a uniform field of view over 200 by 200 μm^2^ and 2 Hz frame rate, we mapped fat storage in free-moving *C. elegans* and live embryos.

## 1. Introduction

Recently developed mid-infrared photothermal (MIP) microscopy, also called optical photothermal IR (O-PTIR) microscopy, has enabled super-resolution IR imaging in the far-field (*1*–*3*). MIP microscopy employs a visible beam to probe a localized and transient temperature rise induced by a nanosecond-pulsed IR excitation. Such local temperature modulation introduces thermal expansion and refractive index alteration. Those changes are collectively revealed by detecting the scattering intensity modulation of a probe beam at visible wavelength, enabling IR spectroscopic imaging at the submicron scale (*4*, *5*). Moreover, the indirect measurement by a visible beam bypasses the water absorption issue and allows photothermal IR imaging of living systems. Since the first high-quality live cell imaging demonstration in 2016 (*4*), this method is quickly expanded with various innovations including fluorescence detection (*6*–*8*), optical phase detection (*9*–*12*), photoacoustic detection (*13*),and integration with computational tomography (*14*, *15*). MIP microscopy has been commercialized into a product termed MiRage, and has enabled very broad applications, as summarized in recent reviews (*1*, *3*).

Optically detected photothermal imaging measures a small modulation on a large background (*16*). Generally, such a task is achieved through heterodyne detection at the IR pulse repetition rate via a lock-in amplifier. Due to the nature of the photothermal process, the demodulation frequency is below megahertz, where large laser noise exists (*17*). To beat the laser noise, a pixel acquisition time of a few milliseconds is required (*2*, *4*, *18*).Spatially multiplexed photothermal detection using a CMOS camera much improved the detection efficiency. In such full-field MIP systems, photothermal contrast is acquired by subtracting the camera-captured frames between IR on and IR off status (*9*, *14*, *19*).However, a common CMOS sensor has a limited photon budget on the level of tens of ke^-^. Consequently, frame averaging is mandatory to resolve the subtle modulation depth (*12*, *19*, *20*). Furthermore, excitation fluence of weakly focused IR significantly diminishes with the field of view. Consequently, high energy mid-infrared laser source must be used for compensation. Collectively, the imaging speed of current infrared photothermal microscope is limited to seconds or minutes per frame for biological specimens, which is insufficient to capture quick dynamics inside living systems or large area mapping with a high throughput. Here, we report a single pulse laser-scan MIP microscope that allows high-sensitivity and high-speed imaging till video rate.

To perform scanning-based imaging at video rate, a sub-microsecond system response at each pixel is required (*21*). Yet, unlike coherent Raman microscopy, the last modulation rate is limited to 1 MHz or lower to avoid thermal accumulation in photothermal microscopy. In such a scenario, the photothermal contrast at each pixel needs to be extracted within a single IR excitation period, in which case lock-in filtering fails to pick the signal from a wideband noise background (*22*). We overcome this difficulty by substituting the lock-in based narrowband detection with a wideband amplifier and a megahertz digitizer. Using this method, the photothermal modulation induced by a single IR pulse can be resolved in the time domain (*17*). With improved system response, a faster imaging scheme rather than sample scanning is needed to match the expected pixel dwell time of microseconds. To address this issue, we employ two sets of galvo for synchronized scanning of both the mid-infrared and the probe beams and achieve a line rate over 2 kHz. These synergistic innovations allow, for the first time, microsecond-scale acquisition of photothermal signal from a single infrared pulse at each pixel. The system provides video rate (25 Hz) imaging (150×100 pixels) of chemical dynamics in a living cell. Moreover, synchronized scanning of IR and visible beam allows uniform illumination of a large field of view (over 400 μm) for high throughput chemical scrutinization. With such capacity, we captured fast lipid dynamics inside a living fungal cell. Video rate imaging further allowed spectroscopic decomposition of a single cell wall. Compared with previous lock-in based, sample-scanning MIP microscope (*4*, *5*, *18*), our new system increases the speed by three orders of magnitude (from millisecond per pixel to microsecond per pixel). Broad applications to a wide range of living systems are demonstrated. Results are detailed below.

## 2. Result

### 2.1 Video rate MIP microscope

As depicted in **Fig.1(a)**, our video-rate MIP imaging system is built on an inverted microscope frame. The IR excitation beam is provided by a pulsed quantum cascade laser (MIRcat 2400, Daylight Solution, US). The IR repetition rate is controlled externally between 500 kHz to 1 MHz, with a duty cycle of less than 30%. The probe beam is provided by a continuous-wave laser with a center wavelength at 532 nm (Samba, Cobolt, Sweden). The probe beam is rapidly scanned by an X-Y galvo mirror with the highest resonant frequency at 3 kHz. The scanned probe beam is conjugated to the back pupil of the objective lens with telecentric relay optics and focused into a sample through a water immersion objective lens with 1.2 NA. The IR excitation is scanned by another pair of XY galvo mirrors. A reflective conjugation using a pair of concave mirrors relays the IR scanning to the back pupil of a reflective objective with 0.5 NA. The design of all-reflective scanning and conjugation avoids the chromatic aberration to meet the requirement for IR spectroscopic imaging. The focus of the IR beam is aligned to overlap with the visible focus before imaging. During the imaging process, the IR focal spot is synchronously scanned with the visible probe, which maintains uniform excitation and probing in a large field of view. The two pairs of galvo are synchronously scanned with an angle scaling factor calculated by the objective focal length and calibrated at the beginning of the experiment.

**Figure 1.**
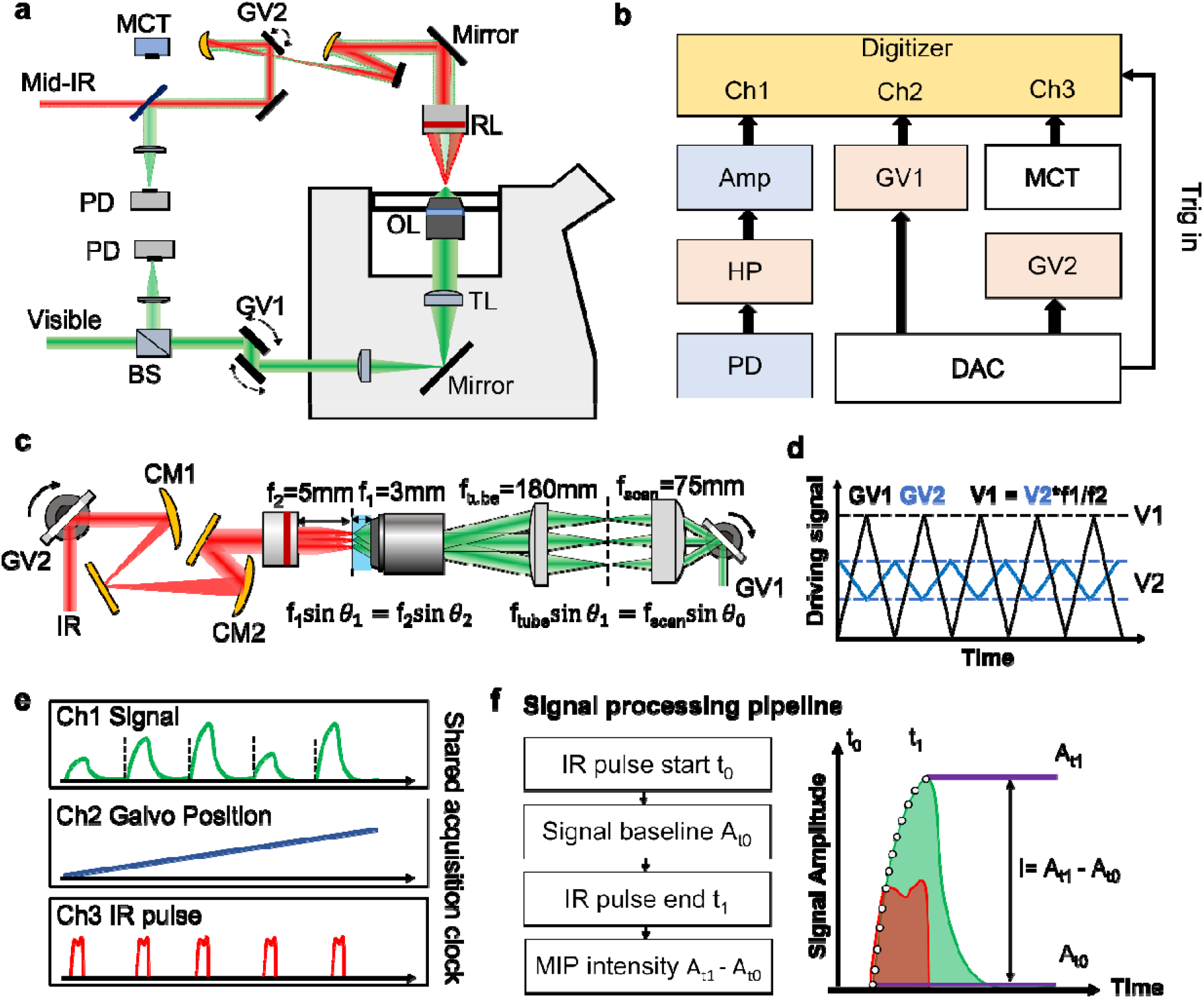
A sync-scan mid-infrared photothermal microscope with single pulse digitization at each pixel. **(a)** Setup. OL: objective lens (60X 1.2NA, UPLSAPO60, Olympus, US), RL: reflective objective lens (40X, 0.5NA, LMM40X-P01, Thorlabs, US), TL: tube lens (f=180mm, Thorlabs, US), SL: scan lens(f=50mm, Thorlabs, US), GV: galvo mirrors (Saturn 1B, ScannerMax, US), PD: photodiode (DET100A, Thorlabs, US), BS: 50/50 beam splitter, MCT: Mercury-Cadmium-Telluride (PVM-10.6, VIGO System, US) detector. **(b)** Control diagram. Digitizer (50 MS/s, Gage Applied, US). Amp: pre-amplifier (SA-230F5, NF, Japan), HP: high pass filter (>0.23 M Hz, ZFHP-0R23, Mini-circuits, US), DAC: digital to analog converter (NI-6363, National instrument, US). The DAC provides the trigger signal for the two galvo scanners and digitizer. The PD and MCT detects the photothermal signal and the IR pulse intensity,respectively. **(c)** Scheme for synchronized scanning of IR excitation and visible probe beams. **(d)** Driver control for galvo synchronization. **(e)** Synchronized recording of MIP intensity, galvo position, and IR pulse intensity at the digitizer. **(f)** Signal processing at each pixel. The IR heating start is monitored by MCT and denoted as t_0_, the corresponding signal amplitude is baseline A_t0_. The peak of temperature rising and photothermal signal A_t1_ arrive at the IR pulse end t_1_. The MIP intensity is a subtraction of peak amplitude A_t1_ by baseline A_t0_.

The probe beam carrying the modulation signals are reversely scanned by the galvo and detected in either forward or backward directions by two silicon photodiodes connected with a preamplifier and filtering circuit. The amplified signal is directly sent to a high-speed digitizer with a sampling rate of 50 million samples per second. The acquisition and control diagram are shown in **Fig.1(b)**. The detailed illustration of synchronized scanning of two counter propagating beam is shown in **Fig.1(c)**. Two pairs of galvo mirrors are controlled with analog driving voltage. The analog driving signals are scaled with corresponding scanning angle given by the focal length and relay optics as shown in **Fig.1(d)**.

During the imaging process, the digitizer synchronously records the MIP signal, galvo position feedback and IR excitation trigger line by line as depicted as **Fig.1(e)**. The single pulse contrast at each pixel is then extracted with the data processing method shown in **Fig.1(f)**. The contrast is a subtraction of the signal amplitude before IR pulse heating (t_0_) from that after IR pulse heating (t_1_). With acquired MIP intensity of each pixel, the image is then reconstructed according to the scanning trajectory revealed by the galvo sensor.

### 2.2 High signal to noise ratio over a large field of view

We used PMMA particles with 500-nm diameter to evaluate the performance of our single-pulse MIP microscope. The PMMA particles in solution form were first diluted with deionized water. One droplet of the solution was then spread on the surface of a calcium fluoride CaF_2_ substrate with 0.2 mm thickness for imaging. The photothermal signal is detected from backward scattering. The QCL laser is at 500 kHz with a 400 ns pulse width. The laser scan imaging is performed with a pixel dwell time of 2 μs, corresponding to single pulse per pixel and video rate for an image of 150×100 pixels. By tuning the IR excitation to 1729 cm^-1^, corresponding to the absorption peak of the C=O bond of PMMA, the single-pulse photothermal signal from a single particle can be clearly resolved by our wideband detection system as shown in **Fig.2(a)**. We further process the signal and extract the modulation amplitude as the contrast of each pixel as illustrated in **Fig.2(b)**. The raster-scanned photothermal image is then reconstructed as shown as **Fig.2(c)**. A signal-to-noise ratio of 86 was successfully obtained for a single 500-nm diameter PMMA particle.

**Figure 2.**
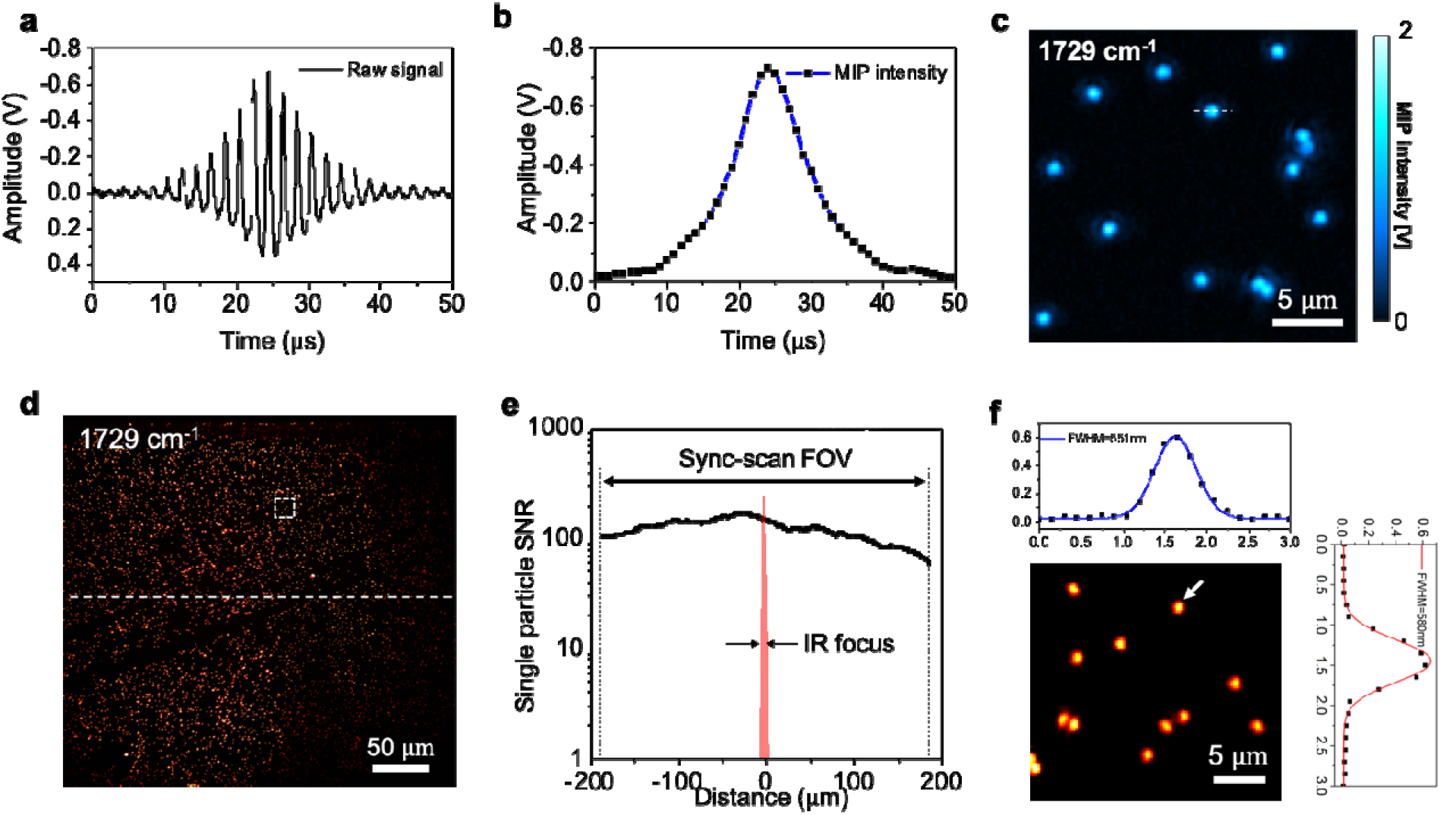
Single-pulse mid-infrared photothermal imaging of 500-nm PMMA particles. **(a)** Raw photothermal signal acquired during laser scanning across a single PMMA particle. **(b)** Extracted MIP contrast from (a). **(c)** Reconstructed photothermal image of 500 nm-diameter PMMA particles generated with a single IR pulse per pixel. IR excitation at 1729 cm^-1^ according to the C=O bond. Indicated dash line corresponds to the signal shown in (a) and (b). Scale bar = 5 m. **(d)** Single pulse imaging of 500-nm diameter PMMA particles in a large FOV. Scale bar = 50 m. **(e)** Single particle SNR evaluated in the whole field of view. **(f)** Zoom-in of the area indicated in (d). The profile is for the particle indicated by an arrow in (f). Calculated spatial resolutions are 231-nm in X-axis and 293-nm in Y-axis, respectively. Scale bar = 5 m.

In addition to video-rate MIP imaging, with synchronously scanned IR and visible lasers, a uniform imaging field of view (FOV) over 400 μm can be maintained without compromising the sensitivity. Here, we show single-pulse imaging of 500-nm PMMA particles in a large area as **Fig.2(d)**. The imaging size is set to 2000×2000 with a pixel size of 200 nm. The chemical contrast is shown by tuning the IR excitation at 1729 cm^-1^. The uniformity is evaluated by single particle SNR across the field of view as shown in **Fig.2(e)**. Leveraging the synchronized scanning scheme, high SNR of ~100 is maintained over 400 μm, which is about 80 times wider than the IR focal spot. The zoom-in image in shown in **Fig.2(f)**. The spatial resolution of 231 nm is calculated from deconvolution of bead diameter from the full width at half maximum. In summary, the laser scanning MIP microscope supports submicron resolution imaging over 400 μm FOV without need of sample movement, which significantly facilitates applications that require high throughput.

### 2.3 Video-rate MIP imaging reveals organelle activity in living fungal cells

Resolving organelle activity in a highly dynamic system requires high detection sensitivity and speed (*23*). Leveraging the large IR absorption cross-section and fast scanning scheme, we demonstrate chemical imaging of small organelle dynamics inside living cells using the single-pulse MIP microscope. Specifically, we acquired the fast lipid dynamics inside fungal cell *C. albicans* at the speed of 20 Hz (Movie S1). The lipids in fungi serve important functions including energy storage, membrane construction, and precursor synthesis (*24*). Here, by tuning the IR excitation to 1740 cm^-1^ corresponding to the absorption peak of the C=O bond in esterified lipid, the individual lipids droplets (LD) can be specifically imaged.

Interestingly, two types of lipid droplets are differentiated based on their distinct dynamics. A larger LD is localized in the cytoplasm with a relatively static position. On the contrary, a fast-moving LD is seen in the vacuole of the cell. The vacuole inside fungal cells plays an important role of LD biogenesis and degradation. The significant lipid motility inside a living fungal vacuole was revealed, as shown in **Fig.3(a)**. With video-rate imaging speed, those fast dynamics can be resolved without distortion. The time lapse projection on the indicated line in **Fig.3(a)** shows that the lipid droplet moved around the vacuole as shown in **Fig.3(b)**. The constant signal level from the cell wall indicates no photobleaching. The time lapse of lipid dynamics within 8 seconds is shown as a color-coded image in **Fig.3(c)** depicting the trajectory of the LD. Collectively, by resolving the fast organelle dynamics, our method can be used to study LD internalization or degeneration in response to nutritional environment alternations, as well as cellular response to a treatment.

**Figure 3.**
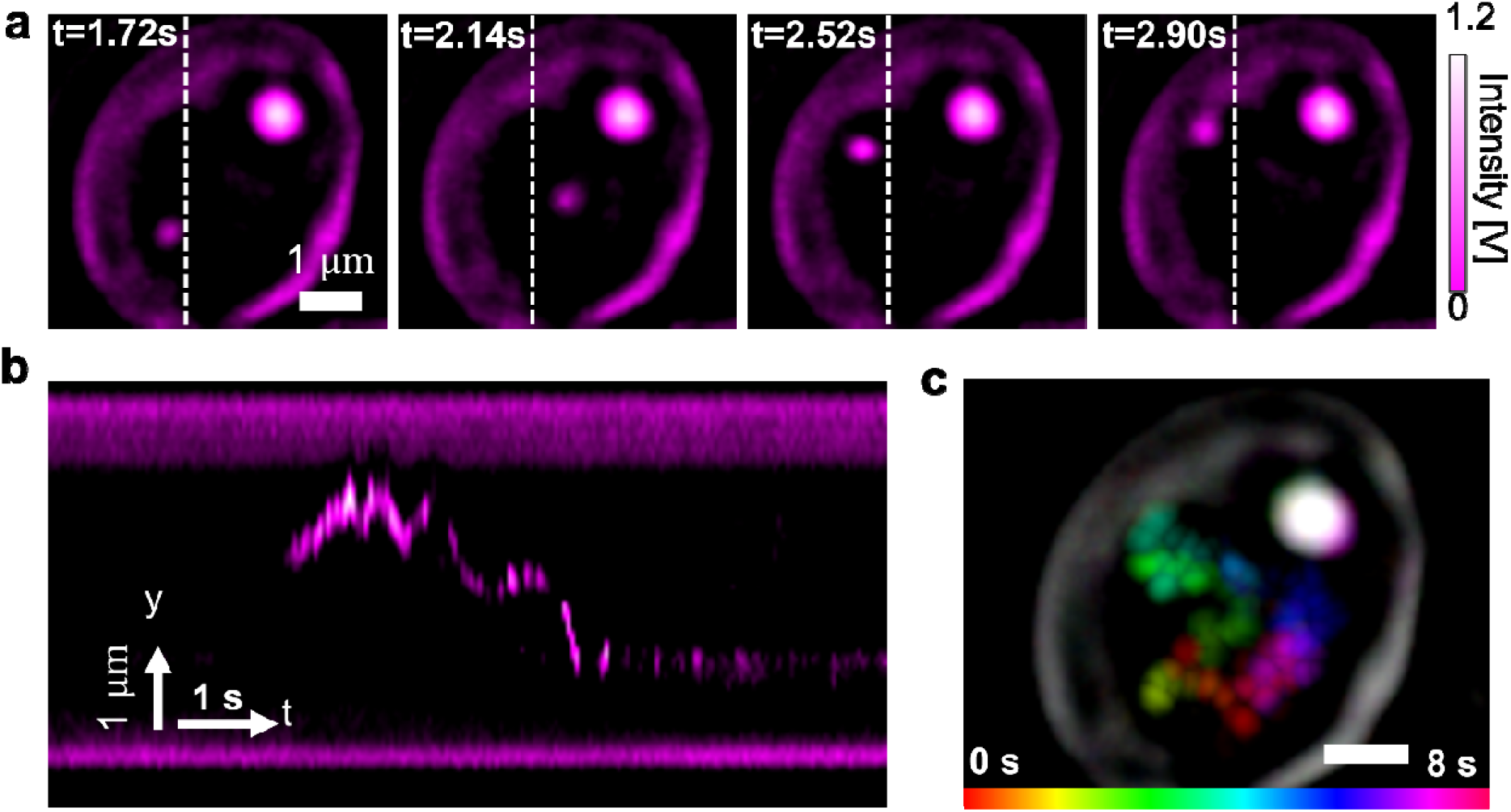
Video-rate MIP imaging reveals fast lipid dynamics inside a fungal vacuole. **(a)** Lipid movement was recorded at different time points. IR excitation tuned to 1740 cm-1 according to the absorption peak of lipids. Scale bar = 1 m. **(b)** Y-t view of the line indicated in a. **(c)** Temporal color-coded image showing the dynamics of the lipid droplets. The static features such as the cell wall show a white color. Scale bar = 1 m.

### 2.4 Spectroscopic MIP imaging reveals layered composition of a fungal cell wall

The fungal cell wall is majorly composed of layered polysaccharides (mannans, glucan, chitin). Imaging the cell wall without labeling remains difficult due to its extreme thin thickness of 100 to 300 nm (*25*). Moreover, fast imaging speed is required to avoid the distortion caused by sample movement. With the chemical specificity and submicron resolution, our single-pulse MIP microscope allows visualization of such structure in a living cell. By tuning the IR excitation to 1740 cm^-1^ that corresponds to the absorption of C=O bond in triglyceride, lipid droplets inside the cells are seen clearly. Strikingly, by tuning the IR excitation to 1070 cm^-1^ that corresponds to the absorption of C-O bond in carbohydrate, the cell wall of individual fungi cells is clearly resolved, as shown in **Fig. 4(a)**. The cell wall at indicated dash line in Fig.4(a) has the FWHM of 326 nm, corresponding to a thickness of 239 nm after deconvolution with PSF.

**Figure 4.**
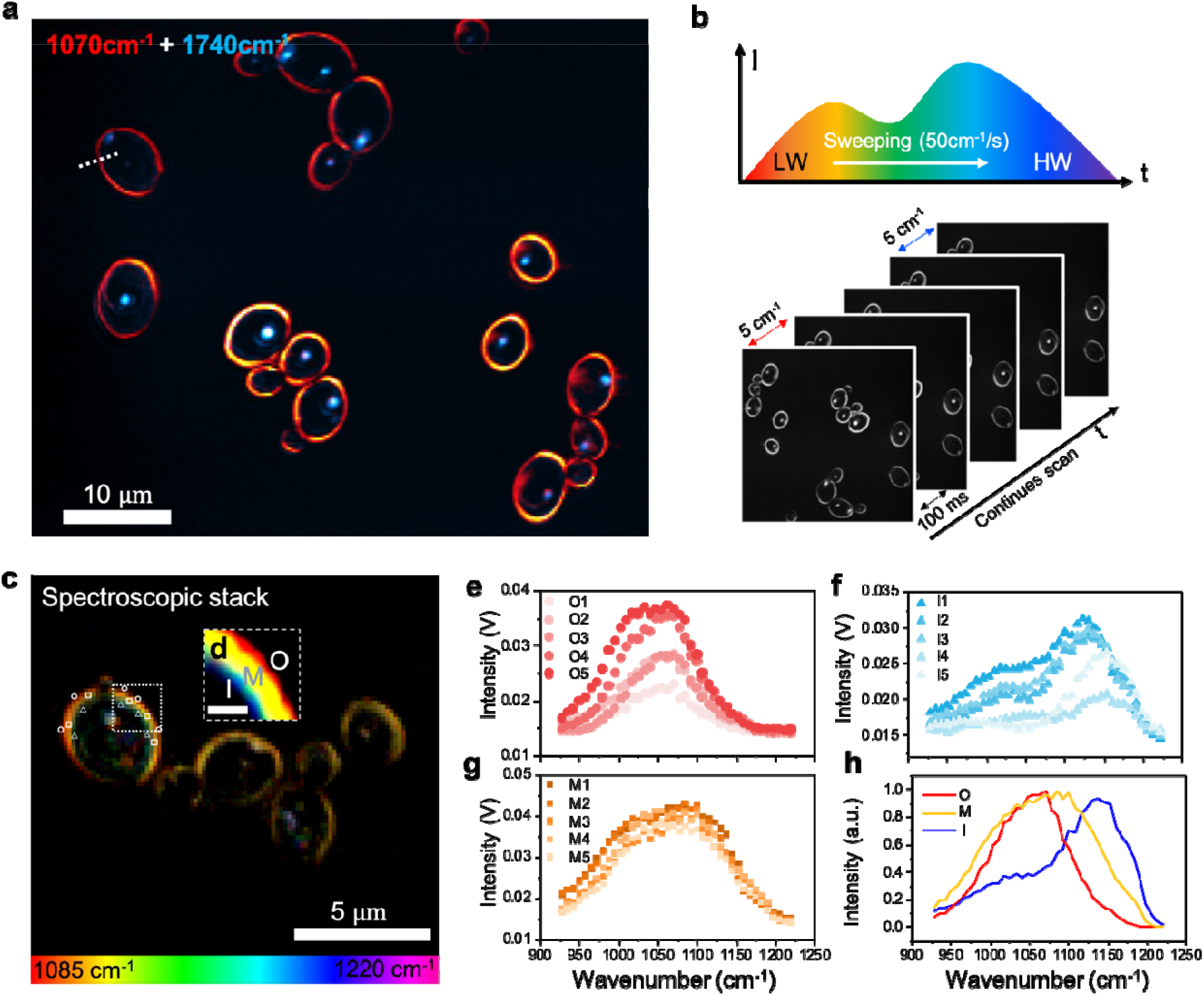
Video-rate MIP enables hyperspectral imaging of fungal cell wall. **(a)** Multi-color images of fungal cells. IR at 1070cm^-1^ excites the polysaccharides components inside the cell, where the cell wall gives a strong signal. **(b)** Hyperspectral imaging scheme. The QCL laser sweeps during the acquisition of MIP videos. Trigger from laser synchronized the frame number with wavenumbers. **(c)** Fast hyperspectral imaging of fungal cell wall with the excitation from 1085 cm^-1^ to 1220 cm^-1^. The entire stack of 60 frames took 6 seconds. Scale bar = 5 m. **(d)** Image deconvolution of indicated area in (c). A three-layer feature can be resolved from the cell wall, indicating a nanoscale compositional difference. O: outer. M: middle. I: inner. Scale bar = 500 nm. **(e, f, g)** Spectra extracted at indicated pixels in (c). (h) Averaged spectra of the three-layer features indicated in (d).

Next, leveraging the improved imaging throughput and the fast-tuning function of the QCL laser, we extended the system’s ability for high-speed mid-infrared spectroscopic imaging as shown in **Fig. 4(b)**. We focused the fungal cell wall and resolved its layered cell wall structure based on composition difference. This is the first report of such fine structure observed chemically in living cells with a far-field optical microscope.

To provide molecular insights into the cell wall composition, spectroscopic imaging was performed by sweeping QCL laser from 900 cm^-1^ to 1200 cm^-1^. We performed imaging at speed of 10 frames per second and swept the laser wavelength at the speed of 50 cm^-1^, offering an effective spectral resolution at 5 cm^-1^ (see Movie S2). The sum-up intensity of the hyperspectral stack is shown in **Fig. 4(c)**. The IR spectrum in the C-O group region can be clearly resolved at each pixel. In the color-coded hyperspectral image, an obvious layered structure of the cell wall can be observed due to the spectrum difference. Image deconvolution of indicated area in **Fig.4(c)** is shown in **Fig.4(d)**, where a distinctive three-layer structure model was seen. This result indicates a higher spatial resolution than single color MIP. The structure below the diffraction limit can be revealed through spectral discrimination. Spectroscopic imaging has been used for improved the spatial resolution of single molecule localization microscopy by using dyes having different excitation and emission spectra (*26*, *27*). Here, we show that spectroscopic MIP imaging could resolve sub-diffraction objects based on their distinct absorption spectra. To further explore the chemical composition, we extract the spectrum of indicated pixels, as shown in **Fig.4(e, f, g)**. The outer layer shows a sharp peak located in 1050 cm^-1^, majorly contributed by mannan (*28*). In the middle of the cell wall, the peak becomes broad and blue-shifted due to the mix of multiple glucan and chitin, which displays a central peak at 1080 cm^-1^. The inner layer is majorly contributed by the membrane which given an absorption peaked at the higher wavenumber of 1150 cm^-1^. The comparison of the three-layer spectra is shown in **Fig.4(h)**. The fungal cell wall plays an important role for maintain cell function, and proliferation (*25*). It is also the most common target of antifungal drugs for pathogenetic yeast. Direct investigation of its chemical composition and morphology change during cell-drug interaction can fuel up the development of new treatment methods.

### 2.5 Large-area single-pulse MIP imaging of chemical dynamics in living systems

To demonstrate the imaging capability on more complex living systems, we further apply the single-pulse MIP system to acquire the dynamics of cancer cells and multicellular organism *C. elegans*. OVCAR-5 ovarian cancer cells were cultured and imaged with living cell medium as buffer. By tuning the excitation into 1553 cm^-1^ according to the absorption peak of Amide-II band, the protein map was required and shown in **Fig.5(a)**. With a pixel dwell time of 2 μs, a frame rate of 8 Hz was selected to map the whole cell body with fine spatial resolution. With MIP signals from proteins, the dynamics of mitochondria and nucleus can be clearly resolved (Movie S3). The movements of indicated area at different time points are shown in **Fig.5(b)**. By tuning the excitation to 1740 cm^-1^ according to the peak of C=O bond in ester, the lipid droplets inside the cell are clearly resolved as shown in **Fig.5(c)**. Unlike the lipid dynamics shown in fungal cells, the lipid droplets in cancer cells show directional transportation (Movie S4), as indicated in **Fig.5(d)**. It is known the lipid droplets participate cell cellular metabolism, coordinates between different organelles as a hub. By resolving the dynamics of such small organelles, our method provides a new scheme for investigating metabolic activity in living conditions.

**Figure 5.**
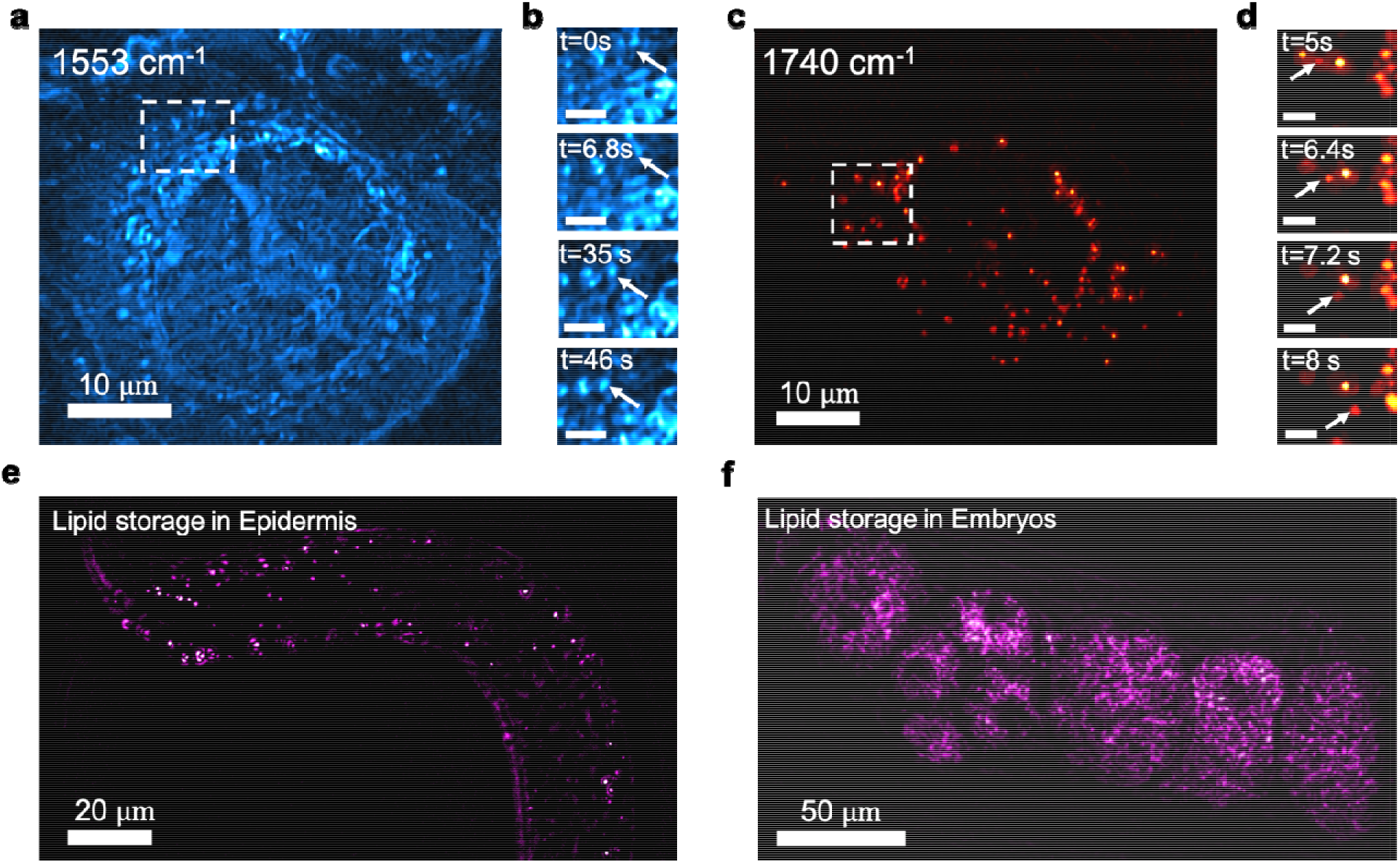
Single-pulse MIP imaging of living cancer cells and *C. elegans*. (a) Protein dynamics in OVCAR-5 with IR excitation at 1553 cm^-1^. Scale bar = 10 m. (b) Intracellular movements at different time point. Scale bar = 2 m. (c) Lipid droplets dynamics with IR excitation at 1740 cm^-1^. Scale bar = 10m. (d) Lipid droplet transportation over time. Scale bar = 2 m. (e) Map of lipid storage in an adult C. *elegans*. Scale bar = 20 m. (f) Map of lipid storage in developing embryos inside a worm. Scale bar = 50m.

To demonstrate in *vivo* chemical imaging ability, we performed single pulse MIP imaging of live C. *elegans,* a model animal extensively used for studying metabolism in disease. For C. *elegans,* Current compositional imaging method highly relies on fluorescent dyes and live imaging is majorly performed at the embryos stage. Imaging postembryonic activity is limited to fixed biological samples. Although transgenic fluorescence has been applied, it remains difficult for resolving various chemical composition inside a live worm (*29*). Leveraging the label-free chemical imaging in nature, we applied our system live C. *elelgans* imaging. Our method is applicable for both embryonic and postembryonic cell dynamics. By tuning the excitation to 1740 cm^-1^, the stored lipid in epidermis can be clearly resolved (see Movie S5) as shown in **Fig.5(e)**, which is the main triglyceride depot. Moreover, by imaging its reproductive system, strong signals from lipid are seen inside the developing embryos (Movie S6) as shown in **Fig.5(f)**. During the imaging period, no photodamage was observed on the living system for continuous imaging over minutes. With the capability of in *vivo* imaging on C. *elegans,* we envision that the single-pulse MIP microscope can provide a new way to investigate the molecules involved in life development.

## 3. Discussion

By high-speed digitization and synchronous scan of IR and visible beams, we have pushed the MIP imaging speed to the level of single IR pulse per pixel, enabling video-rate chemical imaging of living dynamics. With much improved throughput, we further extend the system for fast mid-IR photothermal spectroscopic imaging of living cells. For realizing the speed boost, we first employed a lock-in free demodulation method via high-speed digitization. This method improved the system response for single-cycle demodulation and enabled high detection sensitivity at microsecond pixel dwell time. Subsequently, we implemented synchronized scanning of counter-propagated probe and pump beam to enable photothermal imaging with a line-rate over 2 kHz, in a uniform field of view over 400 m and with a spatial resolution down to 220 nm.

The demonstrated single-pulse per pixel detection scheme can be generally adapted for pump-probe signal extraction to push the imaging speed and sensitivity. It especially boosts the sensitivity of low-duty cycle signal detection, where lock-in demodulation of single frequency is not efficient to separate signal from background noise. With improved high detection sensitivity, fast digitization brings the system response to the nanoseconds level without requiring special electronics design (*21*), allowing the pump-probe contrast extraction within a single excitation cycle. To bring the MIP imaging speed to the microsecond scale, laser scan is used instead of the traditional sample scan approach. The innovative synchronized laser scanning of counter-propagated IR and visible beam provides high imaging quality cross the entire field of view, in terms of uniform signal level and spatial resolution.

Notably, mid-infrared spectroscopic imaging requires a long wavelength tuning range over thousands of wavenumbers. Our all-reflective laser scan design fundamentally solves the chromatic aberration issue, providing quantitative spectroscopic imaging results over the whole fingerprint. The proposed design can be extended to traditional mid-infrared spectroscopic imaging (*30*) to realize a high-speed confocal mid-infrared microscope. Our all-reflective design can be widely adapted to any pump-probe microscopes that are affected by chromatic aberration of two beams.

With the improved imaging speed till video rate, we demonstrated mid-infrared scrutinizing of organelle activity inside a living cell, which is beyond the capability of AFM-IR. In addition to dynamic imaging, our system advanced single-color MIP imaging towards fast spectroscopic imaging. Our results of cell wall imaging showed such ability by revealing the chemical compositional difference at the nanometer-scale. Collectively, with the high sensitivity gained from large IR absorption cross-section and the video rate imaging speed gained from single pulse photothermal detection at each pixel, our work opens a new dimension for biology and chemistry research where analytical tools with high spatial resolution and high speed are required.

## 4. Methods and Materials

### 4.1 Mid-infrared spectroscopic imaging

To perform the mid-infrared spectroscopic imaging, the QCL is operated in continues sweeping mode. The data acquisition system is then triggered by the QCL when the sweeping begins. The acquisition ends when the QCL finishes the sweeping. The start and end frames correspond to the setting wavelength. The hyperspectral data is analyzed and displayed with ImageJ, using temporal color code. Deconvolution is performed using ImageJ on the color-coded image.

### 4.2 PMMA particles

The PMMA particles with a nominate diameter of 500□nm in solution form were first diluted with deionized water. Around 2 μL of the solution was then dropped on the surface of a calcium fluoride (CaF_2_) substrate with 0.2□mm thickness for imaging. The photothermal signal was detected from backward scattering.

### 4.3 Candida albicans

*Candida albicans* isolates (strain number 55) were cultured in yeast extract peptone dextrose overnight at 37□°C with 250□r.p.m. shaking. The 500□μl Candida suspension was centrifuged, washed three times with phosphate-buffered saline and diluted in phosphate-buffered saline. Around 15 μL of the solution was then dropped on the surface of a CaF_2_ substrate with 0.2□mm thickness and sandwiched with 0.15 mm thickness coverslip. The photothermal signal was detected from forward scattering.

### 4.4 OVCAR-5 cancer cells

OVCAR-5 ovarian cancer cells were cultured on CaF_2_ substrate for 24 h. The culture medium was substituted by phosphate-buffered saline right before imaging. During the imaging, the cell was sandwiched between CaF_2_ substrate and 0.15 mm thickness coverslip. Spacer with 0.25 mm thickness was used for maintaining the medium buffer. The photothermal signal was detected from forward scattering.

### 4.5 C. elegans strains

*C. elegans* strains, wild-type (N2) and CB1370 [daf-2(e1370)], were obtained from the Caenorhabditis Genetics Center. *C. elegans* strains were cultured on agar plates seeded with OP50 Escherichia coli. For all experiments, the worms were suspended in phosphate-buffered saline. Around 15 μL of the solution was then dropped on the surface of a CaF_2_ substrate with 1□mm thickness and sandwiched with 0.15 mm thickness coverslip right before imaging.

## Supporting information

Supplementary methods and videos are available as supporting information.

## Data availability

The authors declare that all the data that support the findings of this study are available within the paper and Supplementary Information. Additional relevant data are available from the corresponding authors upon reasonable request.

