## Supplemental materials for "Video-rate Mid-infrared Photothermal Imaging by Single Pulse Photothermal Detection per Pixel"

Jiaze Yin et al.

**This PDF file includes:**

Fig. S1. Zemax simulations of the reflective relay system

Fig. S2. Hyperspectral MIP imaging of 500-nm diameter PMMA particles

Fig. S3. System PSF characterization

Fig. S4. Transmission images of the fungal cell during hyperspectral imaging

Legends for movies S1 to S6

**Other Supplementary Materials for this manuscript include the following:**

Movie S1. Video-rate MIP imaging of lipid dynamics inside a living fungal cell

Movie S2. High-speed hyperspectral MIP imaging of fungal cell wall
Movie S3. Single-pulse MIP imaging of protein dynamics in a living cancer cell

Movie S4. Single-pulse MIP imaging of lipid dynamics in a living cancer cell

Movie S5. Single-pulse MIP imaging of lipid storage in free-moving *C. elegans*

Movie S6. Single-pulse MIP imaging of lipid inside developing embryos of *C. elegans*

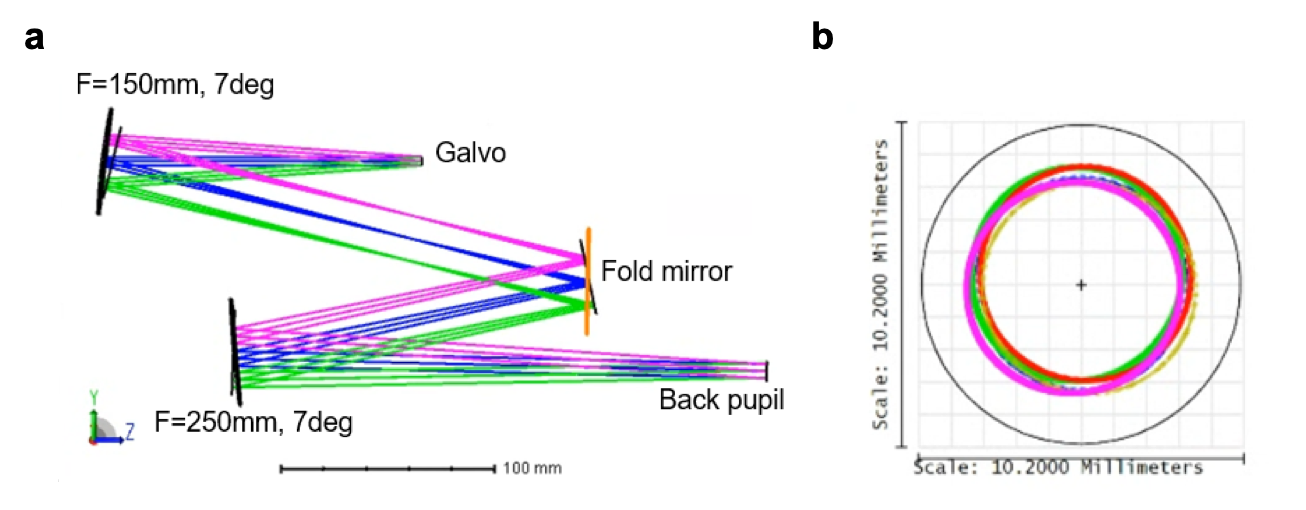

Fig. S1. Zemax simulations of the reflective relay system. (a) Schematic diagram. The beam is simulated with scan angles from -4 degrees to +4 degrees. (b) Ray tracing result at the objective black pupil. The beam spot has a spatial offset of less than 0.6 mm at the maximum scan angle.

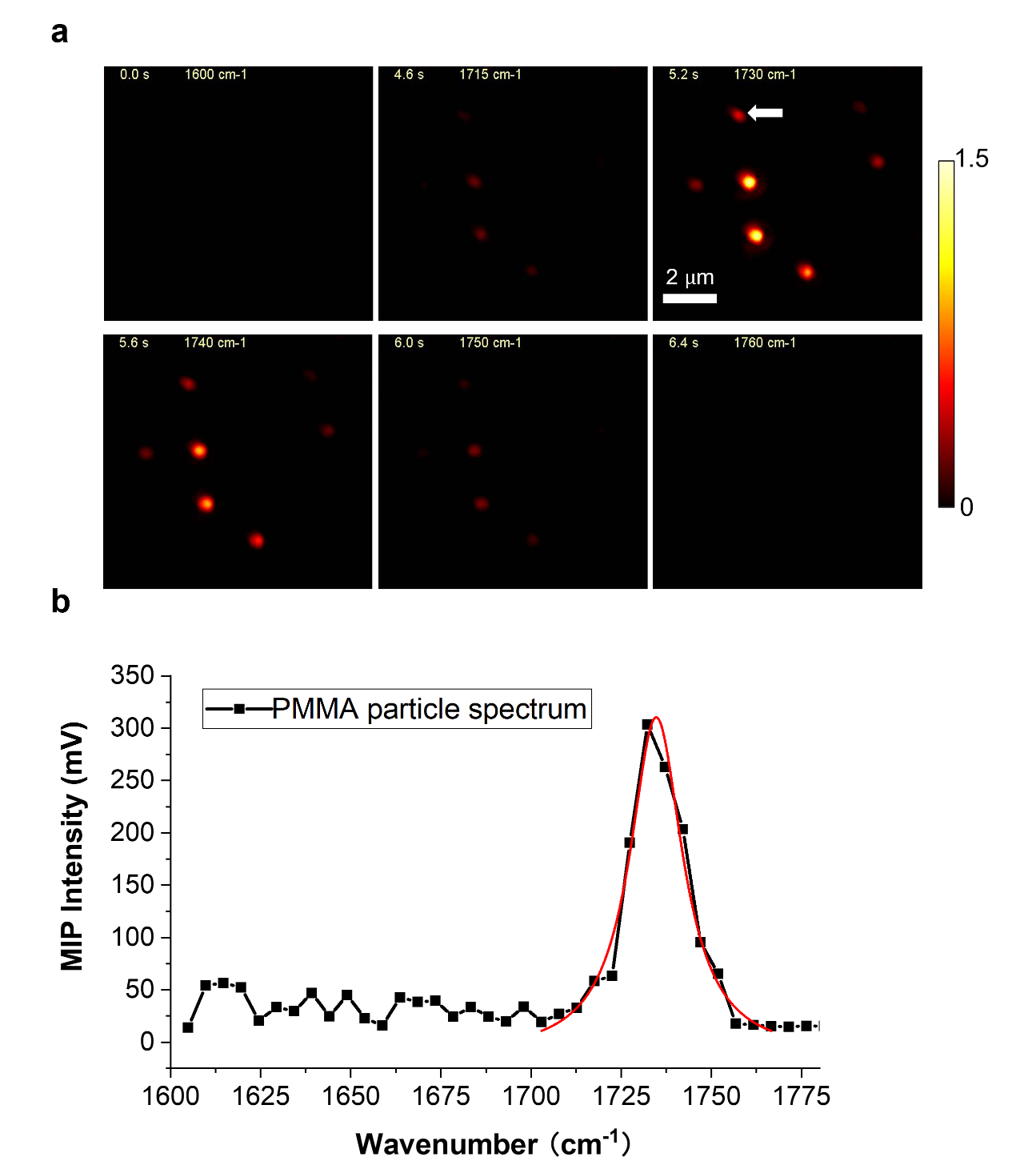

Fig. S2. Hyperspectral MIP imaging of 500-nm diameter PMMA particles. (a) MIP images at different wavelengths. The MIP imaging is performed at 20 Hz with QCL sweeping at 100cm^-1/^s. (b) The spectrum of indicated PMMA particles in (a). The peak of the C=O bond at 1730cm^-1^ can be clearly resolved.

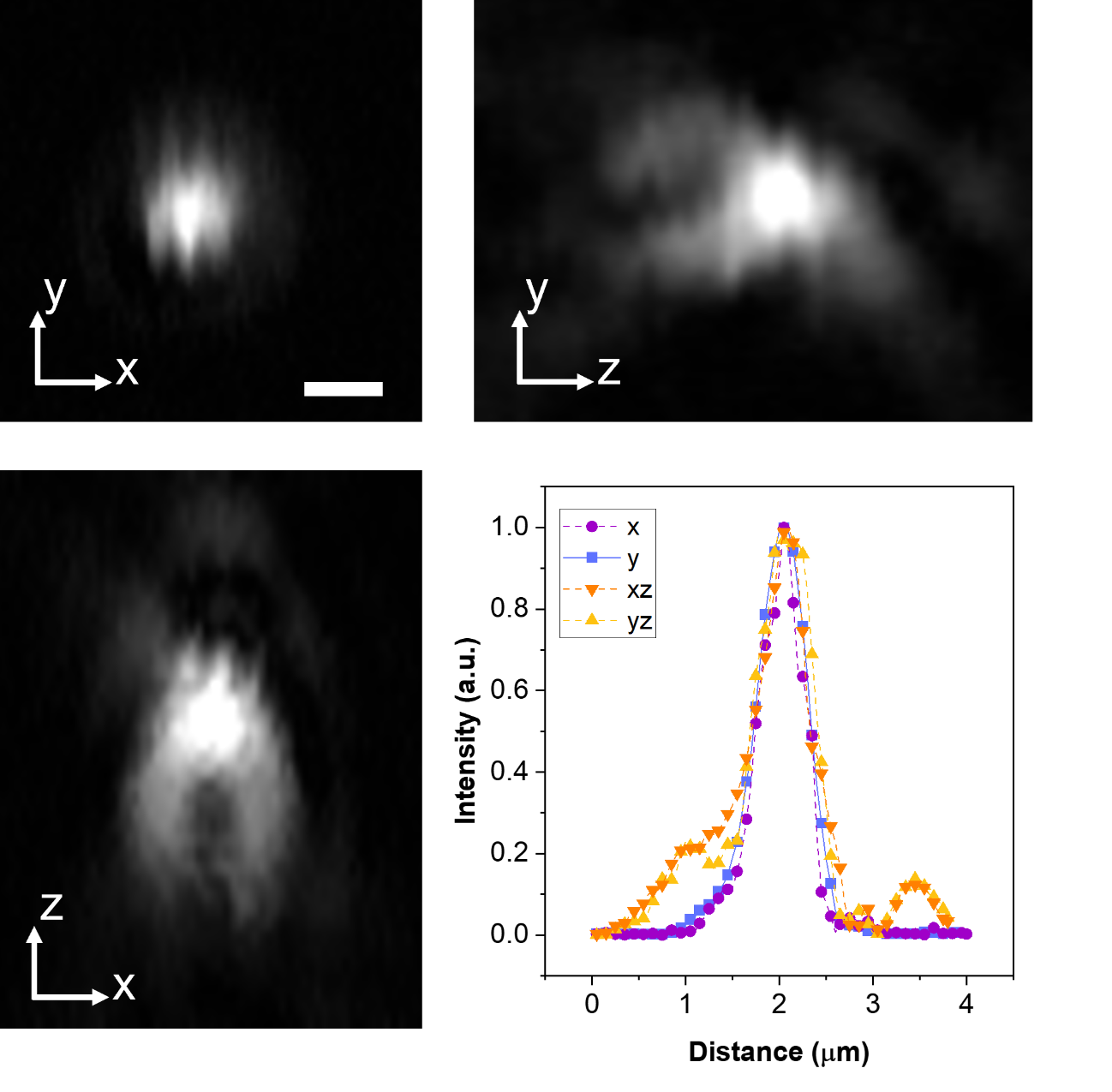

Fig. S3. System PSF characterization. MIP imaging of 500-nm diameter PMMA particles. Projections along xy, xz and zy are shown. The line profiles of the PSF were plotted and fitted along the three principal axes. The FWHM of x, y, xz, yz, respectively, 551 nm, 570 nm, 632 nm, 661 nm, corresponding to a lateral resolution of 231 nm and axial resolution of 386 nm. Scale bar, 1 μm.

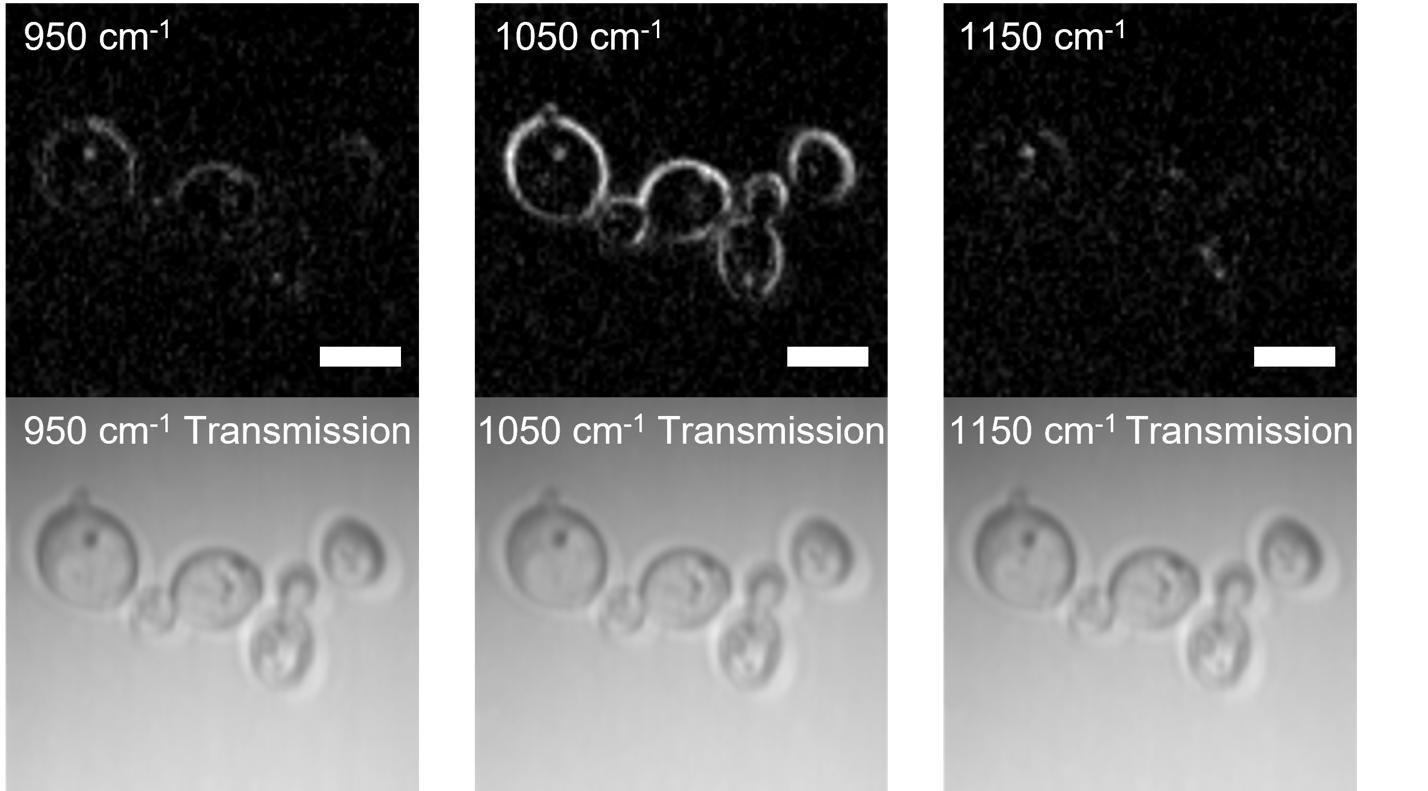

**Fig. S4** **Transmission images of the fungal cell during hyperspectral imaging.** Fungal cells keep the morphology during the hyperspectral acquisition process.

**Movie S1. Video-rate MIP imaging of lipid dynamics inside a living fungal cell.** The movie file visualizes the fast lipid movement at 20 Hz. Each frame with 150 by 100 pixels is acquired with pixel dwell time of 2 $\mu$s. Extra 60 pixels are padded to each line for achieving linear gavlo movement.

**Movie S2. High-speed hyperspectral MIP imaging of fungal cell wall.** The movie file visualizes the hyperspectral MIP imaging process at 10 Hz. The IR source swept wavelength from 900 cm^-1^ to 1200 cm^-1^ at the speed of 50 cm^-1^ per second. Each frame with pixel 200 by 200 pixels is acquired with pixel dwell time of 2 $\mu$s. Total 60 frames acquired provides an effective spectrum resolution of 5 cm^-1^.

**Movie S3. Single-pulse MIP imaging of protein dynamics in a living cancer cell.** The movie file visualizes the protein dynamics in a living OVCAR-5 cancer cell at 8 Hz. Each frame with pixel 250 by 200 pixels is acquired with pixel dwell time of 2 $\mu$s. Extra 50 pixels are padded to each line.

**Movie S4. Single-pulse MIP imaging of lipid dynamics in a living cancer cell.** The movie file visualizes the lipid dynamics in a living OVCAR-5 cancer cell at 8 Hz. Each frame with pixel 250 by 200 pixels is acquired with pixel dwell time of 2 $\mu$s. Extra 50 pixels are padded to each line.

**Movie S5. Single-pulse MIP imaging of lipid storage in free-moving *C. elegans*.** The movie file visualizes the lipid stored in epidermis of adult *C. elegans* at 2 Hz. Each frame with pixel 500 by 500 pixels is acquired with pixel dwell time of 2 $\mu$s.

**Movie S6. Single-pulse MIP imaging of lipid inside developing embryos of *C. elegans*.** The movie file visualizes the lipid stored in developing embryos inside *C. elegans* at 2 Hz. Each frame with pixel 500 by 500 pixels is acquired with pixel dwell time of 2 $\mu$s.
